# Influence of salinity on the thermal tolerance of aquatic organisms

**DOI:** 10.1101/2024.07.11.603038

**Authors:** Luan Farias, Bánk Beszteri, Andrea M. Burfeid Castellanos, Annemie Doliwa, Julian Enss, Christian K. Feld, Daniel Grabner, Kathrin P. Lampert, Serge Mayombo, Sebastian Prati, Christian Schürings, Esther Smollich, Ralf B. Schäfer, Bernd Sures, T.T. Yen Le

**Affiliations:** Department of Aquatic Ecology, Faculty of Biology, University of Duisburg-Essen, Germany; Centre for Water and Environmental Research, University of Duisburg-Essen, Germany; Department of Phycology, Faculty of Biology, University of Duisburg-Essen, Germany; Institute of Zoology, University of Cologne, Germany; Research Center One Health Ruhr of the University Alliance Ruhr, Faculty of Biology, University of Duisburg-Essen, Germany; Ecotoxicology, Research Center One Health Ruhr of the University Alliance Ruhr, Faculty of Biology, University of Duisburg-Essen, Germany

**Keywords:** Co-tolerance, Thermal performance, Osmotic stress, Multiple stressors, Algae, Invertebrates, Fish, Meta-analysis, Global changes

## Abstract

Aquatic organisms are challenged by changes in the external environment, such as temperature and salinity fluctuations. The response of an organism to temperature changes can be modified by salinity, thus pointing at the potential interaction of both variables. In the present study, we tested this assumption for freshwater, brackish, and marine organisms, including algae, macrophytes, heterotrophic protists, parasites, invertebrates, and fish. We reviewed the existing body of literature on potential interactions between temperature and salinity and performed a meta-analysis that compared the thermal tolerance (characterized by the temperature optima, lower and upper temperature limits, and thermal breadths). The final database includes 90 relevant publications (algae: 15; heterotrophic protists: 1; invertebrates: 43; and fish: 31). Relevant publications for microphytes and parasites were not available. Overall, our results show that decreasing salinity significantly increased the lower temperature limits and decreased the upper temperature limits irrespective of the organism groups. These findings mainly reflect the response to salinity changes in brackish and marine systems that dominate our database. Although the number of studies on freshwater species was limited, they showed negative, although statistically nonsignificant, effects of an increased salinity on the thermal tolerance of these species (i.e. increased lower limits and decreased upper limits). In addition, our meta-analysis shows nonsignificant differences in the responsiveness of thermal tolerance to salinity changes among different groups of organisms, but the sensitivity of thermal tolerance to salinity changes generally followed the order: algae > invertebrates > fish. Facing the impact of climate change, our findings point at adverse effects of salinity changes on the temperature tolerance of aquatic organisms. Further studies that investigate the thermal performance of freshwater species at various salinity gradients are required to broaden the evidence for interactions between salinity and temperature tolerance. This also applies to the influence of parasitic infections, which have been found to modulate the temperature tolerance of aquatic invertebrates and fish.

## 1. Introduction

Temperature and salinity are among the most important environmental variables governing the distribution and development of organisms in aquatic ecosystems (Bonacina et al., 2023; Duan et al., 2023). Aquatic systems are subjected to substantial fluctuations of both variables. The underlying drivers of elevated water temperature include climate change, i.e. increases in air temperature and anthropogenic heat emission (Liu et al., 2020). Aquatic ecosystems are additionally challenged by changes in ion concentrations. Salinity increases in freshwater ecosystems, hereafter called salinisation (Reid et al., 2019), are driven by climate change, agriculture (associated with increased surface runoff), mining (associated with the discharge of highly saline water), and road runoff (Canedo-Argüelles et al., 2016, 2013; Feld et al., 2023; Schröder et al., 2015; Hintz et al., 2022). Different patterns of salinity changes in marine ecosystems have been reported. At high latitudes of the oceans, global warming-increased net precipitation dilutes the concentration of ions, contrasting with salinization due to decreased net precipitation and elevated evaporation at low latitudes (Curry et al., 2003; Boyer et al., 2005). Saline rivers, estuaries, and salt-marshes are being diluted, mainly owing to runoff decreases (Zhang et al., 2021).

Generally, responses of organisms to binary stressors can be classified into three types: random co-tolerance (i.e. tolerance to stressors is unrelated), cross-tolerance (or positive co-tolerance), and cross-susceptibility (or negative co-tolerance) (Vinebrooke et al., 2004). Cross-tolerance occurs when the increase of one stressor enhances the tolerance to another (Todgham et al., 2005). This phenomenon might result from triggering similar defence mechanisms or response pathways (Isaza et al., 2021; Korkaric et al., 2015; MacMillan et al., 2009; Todgham and Stillman, 2013; Vergauwen et al., 2013). By contrast, cross-susceptibility occurs when exposure to one stressor reduces the tolerance towards the other stressor. Cross-susceptibility may originate from trade-offs in the ability of organisms to tolerate each stressor (Cuenca-Cambronero et al., 2021; Sinclair et al., 2013; Todgham and Stillman, 2013; Vinebrooke et al., 2004); or shared damage that leads to disproportionately higher effects compared to the individual stressors (Cruz-Loya et al., 2021). For example, there is a trade-off between the energy demands for osmoregulation and the ability to protect the organism against other stressors such as changes in temperature (Vereshchagina et al., 2016).

Changes in temperature and salinity can jointly act on aquatic organisms as thermal and salinity stressors activate the same cellular response, and trigger the expression of the same genes in aquatic organisms (Lockwood et al., 2010; Lockwood and Somero, 2011). For aquatic organisms, osmoregulation is crucial to sustain an osmotic balance between the internal fluids and the external environment. Changes in the concentration of ions in the external environment can increase the costs of osmoregulation. However, our understanding of the response of thermal tolerance to salinity gradients is poor, and it is unknown whether the responses differ systematically between organism groups. In the present study, we compare the thermal tolerance of various groups of eukaryotic aquatic organisms at various salinity levels to identify the type of co-tolerance per organism group. We hypothesize that due to the trade-offs in the energy demand and shared damages, as explained above, the correlation between thermal tolerance and salinity tolerance would be negative, i.e. negative co-tolerance. In other words, we expect salinization would decrease the thermal tolerance of freshwater species (see Le et al., 2023), while salinity dilution would negatively affect the thermal tolerance of marine species. In addition, we hypothesize differences in the co-tolerance among organism groups. Osmoregulatory capacities might be insufficient for microorganisms at low trophic levels, such as algae and fungi, to cope with salinity changes. Therefore, we expect that their thermal tolerance is more responsive to salinity fluctuations.

## 2. Materials and Methods

### 2.1. Database construction

A database of thermal tolerance at various salinity levels was built by retrieving the relevant variables (e.g. optimum temperature, temperature limits, incipient temperature, and thermal breadth) from peer-reviewed publications available through the Web of Science and Scopus. The literature survey was conducted on 18^th^ January 2024 using a combination of various terms describing the responses of aquatic organisms to temperature and salinity changes (Appendix 1: Database construction). After checking for duplicates, the retrieved records were screened following a stepwise approach (Fig. 1). ASRreview abstract and manual abstract screening were to remove irrelevant records (e.g. field or community studies), while manual full text screening was to retrieve relevant records. The machine learning tool ASReview (Active learning for Systematic Reviews) has been demonstrated to be an efficient reviewing tool (van de Schoot et al., 2021; Quan et al., 2024). We used this tool for the first screening step that was followed by manual abstract screening to further remove irrelevant records (e.g. studies on the influence of temperature and/or salinity on another stressor). In the final screening step (i.e. manual full text screening), records are considered relevant when they met the following criteria: 1) Records provide at least one indicator of thermal tolerance (Fig. 2; Table S1, Supporting Information). The records that contain equations for determining at least of the indicators were also considered relevant; 2) Effects of temperature and salinity were simultaneously investigated; 3) Data were reported for at least three salinity levels; 4) Data were obtained from laboratory experiments or modelling research based on laboratory experiments under controlled conditions in which other factors besides temperature and salinity were kept constant among treatments; and 5) Data were based on measured responses of populations (for algae and macrophytes) and individuals (for other organisms) in aquatic life stages, i.e. excluding non-aquatic life stages, where applicable. We focussed on eukaryotic aquatic organisms, covering aquatic algae (green algae, diatoms, and dinoflagellates); heterotrophic protists; submerged or floating macrophytes; parasites; fungi (except food fungi), invertebrates, and fish. We excluded studies in which thermal tolerance was determined under irrelevant environmental conditions, for example, air exposure. A detailed description of the steps for database construction is given in Appendix 1.

**Fig. 1.**
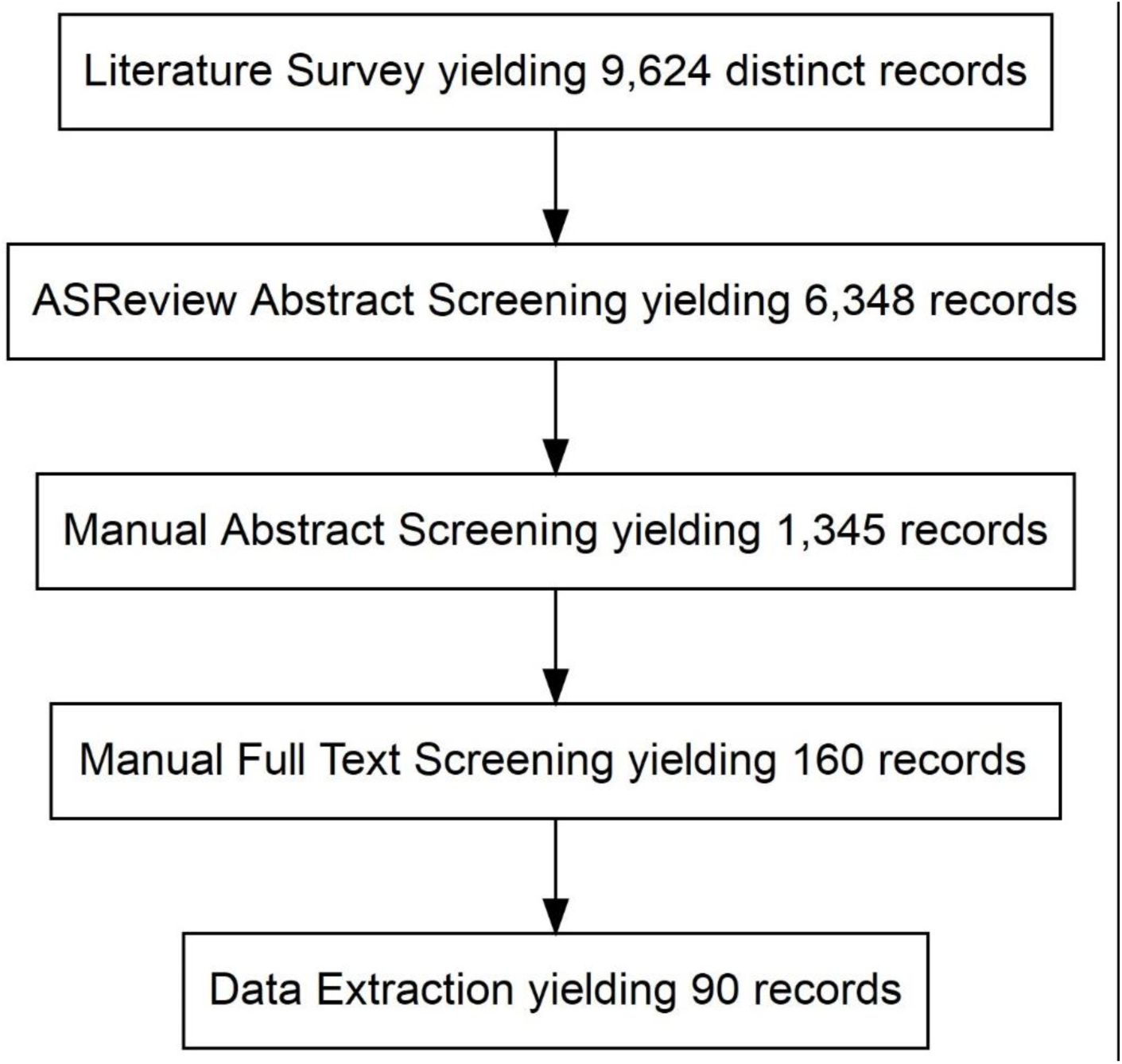
Flow chart of database construction

**Fig. 2.**
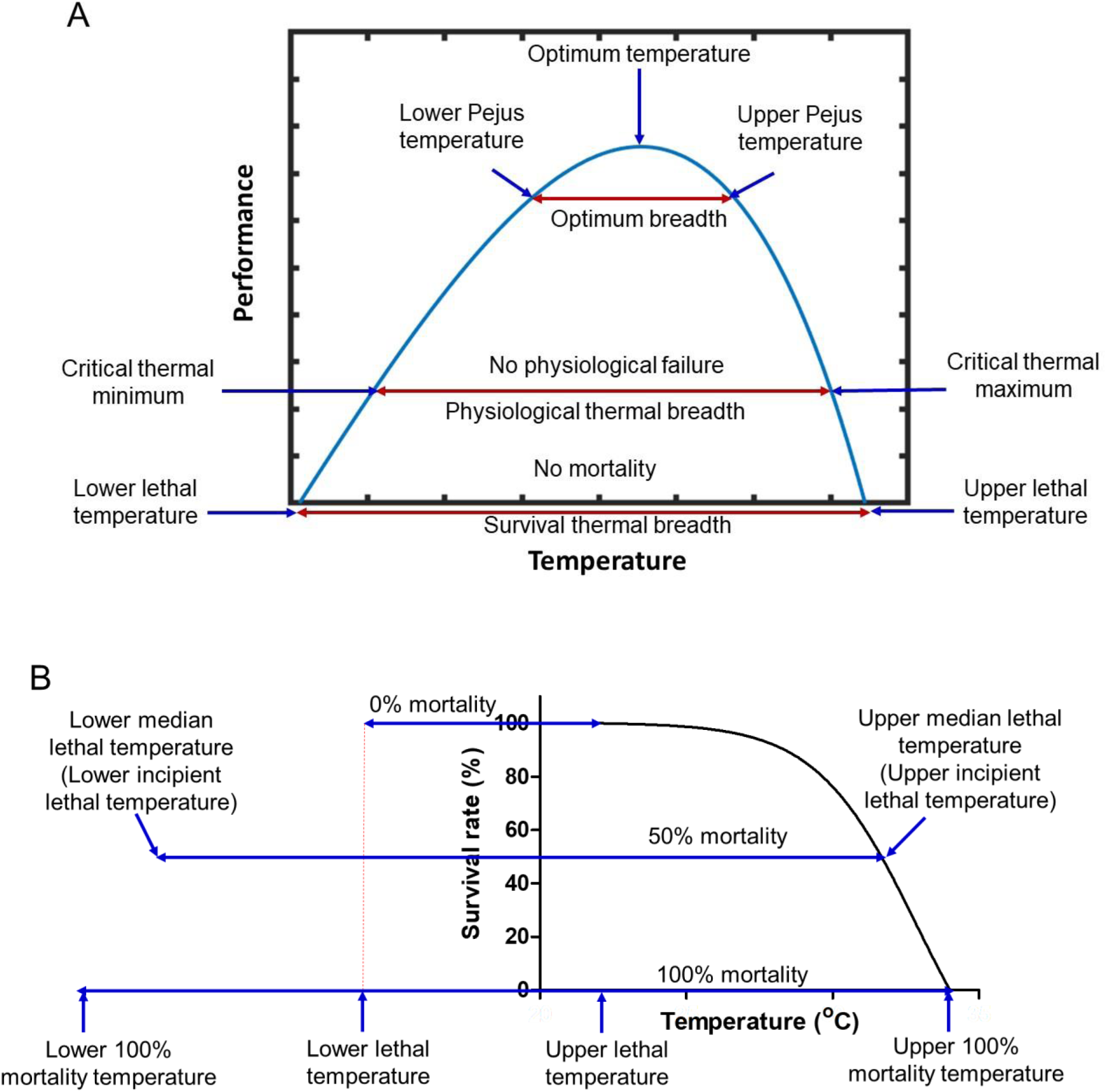
Indicators of thermal tolerance: (A) showing responses to temperature gradients before mortality occurs; and (B) showing change in the survival probability to temperature gradients.

From each relevant record, we extracted the following information, if available, and included them in our thermal tolerance database: species, organism group, habitat, acclimation conditions, test conditions (especially test salinity), indicators of thermal tolerance (indicator, definition, measurements, number of treatments, size of variations, measurement of variations, and number of replicates in each treatment). Data was retrieved from narrative descriptions, tables, figures (using WebPlotDigitizer), and equations in the main text as well as in the Supporting Information. When the experiments were repeated, the mean and its standard deviation were calculated. The final database included 90 publications (organism groups: algae: 15; heterotrophic protists: 1; invertebrates: 43; and fish: 31; habitats: freshwater: 17; brackish: 21; and marine: 53) (Appendix 2: Database). Studies on parasites and macrophytes that met our inclusion criteria were not available. Data on lower and upper ultimate incipient lethal temperatures, as well as on lower and upper ultimate critical thermal minima and maxima were relatively scarce (Table S1, Supporting Information). Therefore, they were omitted from the analyses. Prior to data analysis, the other indicators of thermal tolerance were grouped into six groups: optimum temperature (including optimum temperature and preferred temperature); lower temperature limit (including critical thermal minimum and lower lethal temperature); upper temperature limit (including critical thermal maximum and upper lethal temperature); thermal breadth (including survival and physiological thermal breadth); lower effect temperature (including lower incipient lethal/physiological temperature and lower 100% mortality temperature); and upper effect temperature (including upper incipient lethal/physiological temperature and upper 100% mortality temperature). When both incipient and 100% mortality temperatures were reported from the same study, the incipient temperature was used in the analysis.

### 2.2. Statistical analyses

We aimed to quantify the influence of salinity upon the selected thermal indicators: optimum temperature, lower and upper temperature limits, breadth of the temperature tolerance range, and lower and upper effect temperatures. All measurements of ion concentrations were converted to salinity in ppt. Electrical conductivity was converted to salinity (ppt) using the equation derived by Lewis and Perskin (1981). The effect of salinity on each response was quantified by the slope coefficient of a linear regression model, while the slopes standard error was used to quantify its variation. Since both the predictor (salinity) and the responses (thermal tolerance indicator) were expressed on the same numerical scales across all reviewed studies. We ran weighted random-effects meta-analyses for each thermal indicator as response using the metafor package in R (Viechtbauer, 2010). The weight for each study was assigned based on its number of replicates weight increasing with replicates. Additionally, we considered effects of the following moderators: organism group, habitat, and acclimation conditions (i.e. test organisms were acclimated to test salinity or not). We refrained from using more than one moderator at a time owing to the relatively small sample size. For each response, models with different moderators were compared based on the Akaike Information Criterion (AIC) (Akaike, 1974). We then proceeded to subgroup analyses, applying the same categories to display and examine the differences between these subgroups. Meta-analyses were conducted using RStudio (version 2023.09.1+494, https://github.com/rstudio/rstudio/tree/v2023.09.1+494), and the packages: Tidyverse 2.0.0 (Wickham et al., 2019), lme4 1.1.35.1 (Bates et al., 2015), blme 1.0.5 (Chung et al., 2013), and metafor 4.4.0 (Viechtbauer, 2010). Responsiveness of thermal indicators was classified using the effect sizes of the meta-analyses. Effect sizes were considered significant when the 95% confidence interval excluded zero.

## 3. Results and Discussion

### 3.1. Salinity changes and thermal tolerance

In general, our analysis indicates a positive relationship between salinity changes and the thermal tolerance of aquatic organisms (i.e. salinization extended the thermal tolerance and salinity dilution contracted the thermal tolerance). Increased salinity tends to broaden the thermal tolerance of organisms, while decreased salinity narrows down the thermal tolerance (Table 1). In particular, salinization significantly decreased the lower temperature limit (Fig. 3) and the lower effect temperature (Fig. S1, Supporting Information) as shown by negative pooled effect sizes. Such effects of salinity changes on these two indicators are of a similar magnitude on these indicators as shown by the overlapping effect sizes (Table 1). By contrast, the upper temperature limit (Fig. S2) and the upper effect temperature (Fig. S3, Supporting Information) significantly increased with salinization, as displayed by the positive pooled effect sizes. These two indicators were influenced by salinity changes at the same order of magnitude (Table 1). Moreover, the upper temperature limit was less responsive to salinity gradients than the lower limit (Table 1), which is consistent with the findings of Botella-Cruz, M. et al. (2016). This phenomenon has been defined as the asymmetric response of the thermal tolerance (Botella-Cruz, M. et al., 2016).

**Fig. 3.**
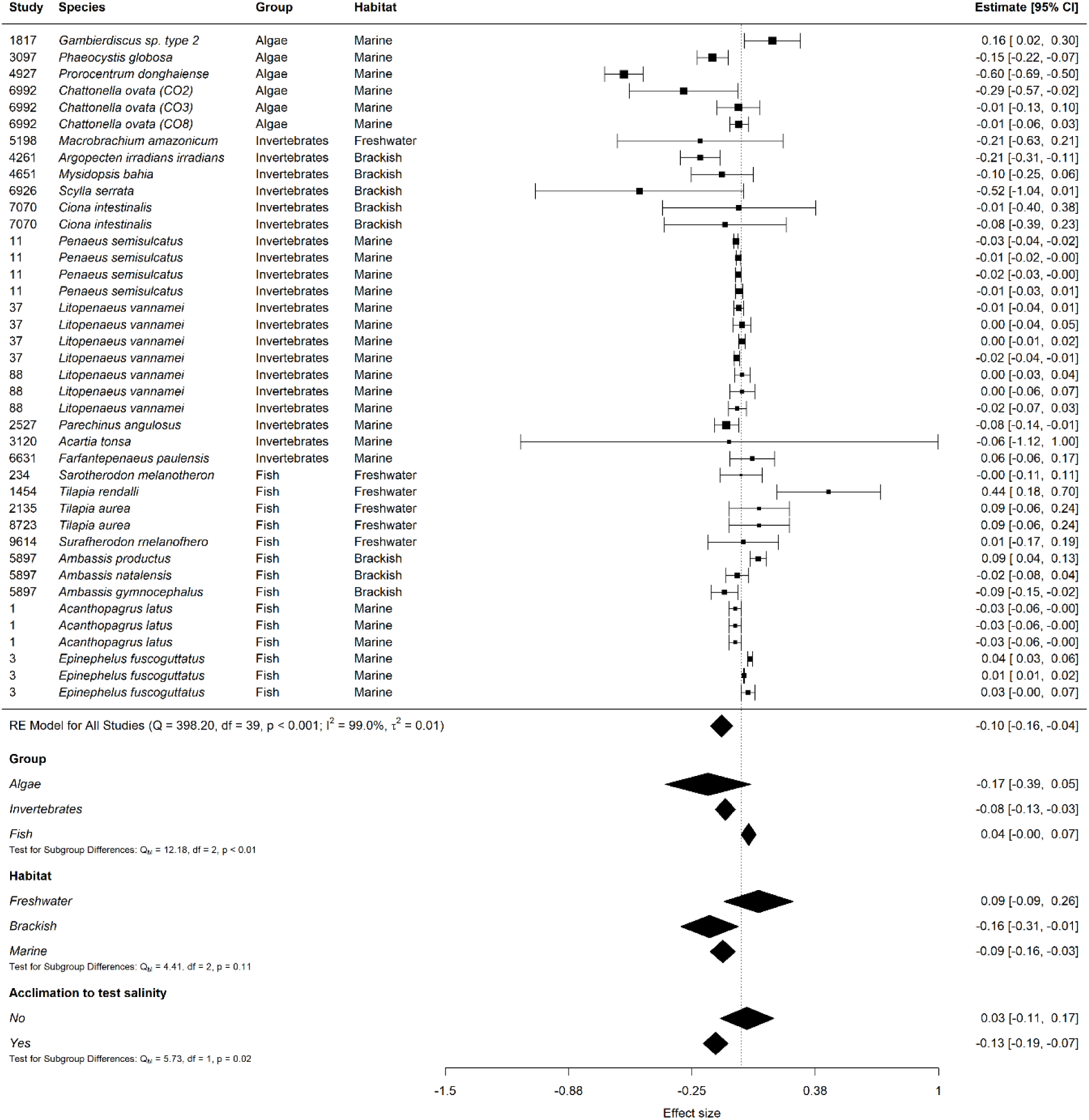
The effect of salinity on the lower temperature limit. Mean (individual and pooled) effect size and 95% confidence interval were estimated from the model. Their values are listed on the right side of the figure. The polygons at the bottom of the plot represent the mean effect size by different subgroups (organism group, habitat, and acclimation to test salinity). Significant effects are defined as the 95% confidence interval not overlapping with zero. Heterogeneity statistics of the model are shown by the values of the Cochran’s Q, its degree of freedom (df), *p*, I^2^, and τ^2^.

**Table 1.** Effect of salinity on the indicators of thermal tolerance. Data availability regarding each response is indicated in terms of both the number of studies and the number of species. Effect sizes, 95% confidence intervals, and p-value (p_mod_) were derived from the meta-analysis for the six thermal indicators. Significant effects are defined as the 95% confidence interval not overlapping with zero. Heterogeneity is expressed by Cochran’s Q and its significance (p_Q_).

| Thermal indicator | Data availability | | Effect size<br>(95% CI) | $p_{\text{mod}}$ | Q | $p_Q$ |
| --- | --- | --- | --- | --- | --- | --- |
|  | Study | Species |  |  |  |  |
| Optimum temperature | 28 | 31 | 0.05<br>(-0.00 – 0.10) | < 0.05 | 1,303,585 | <0.0001 |
| Lower temperature limit | 22 | 26 | -0.10<br>(-0.16 – -0.04) | < 0.001 | 396 | <0.0001 |
| Upper temperature limit | 28 | 35 | 0.05<br>(0.02 – 0.09) | < 0.005 | 9,099 | <0.0001 |
| Thermal breadth | 15 | 19 | 0.11<br>(-0.03 – 0.24) | 0.12 | 215 | <0.0001 |
| Lower effect temperature | 41 | 46 | -0.07<br>(-0.13 – -0.01) | < 0.05 | 3,223 | <0.0001 |
| Upper effect temperature | 46 | 51 | 0.06<br>(0.01 – 0.11) | < 0.05 | 7,611 | <0.0001 |

Brackish (21/90) and marine (53/90) species dominated in our database. This is in line with Velasco et al. (2019) where the majority of experimental studies on multiple-stressor effects also were from brackish and marine species. Hence, the above-mentioned results mainly reflect the response of brackish and marine species (Fig. 3; Figs. S1-S3, Supporting Information). The increased lower temperature limit and the decreased upper temperature limit of brackish and marine species with salinity dilution are consistent with our hypothesis. Salinity decreases might increase energetic costs for adjusting osmoregulation in marine fish and invertebrates (Botella-Cruz, M. et al., 2016; Röthig et al., 2023). Osmoregulation is considered an energetically expensive process as energy is diverted into acclimation and cellular protection (Sokolova et al., 2012). Therefore, salinity dilution may limit the energy available for thermoregulatory mechanisms, such as the production of heat shock proteins, thereby affecting the thermal tolerance (Botella-Cruz, M. et al., 2016; Tomanek et al., 2011). Besides the effects on energy acquisition and allocation, salinity stress might lead to dehydration, thus causing changes in membrane fluidity and consequently decreasing the thermal tolerance (Everatt et al., 2013; Kikawada et al., 2006).

With these results, the thermal breadth is expected to be positively correlated with salinity changes (i.e. extended by salinity increases and contracted by salinity dilution). That was also revealed in our meta-analysis as the average effect size was clearly positive (Table 1; Fig. 4). Notwithstanding, this correlation was not statistically significant, because the confidence interval contained zero (Table 1; Fig. 4). A similar pattern was found for the optimum temperature (Table 1; Fig. S4). Conclusive judgments on the effect of salinity upon the optimum temperature and the thermal breadth could not be obtained as the *p*-values were being or close to significance (Table 1). Our analysis revealed indicator-specific responses of thermal tolerance to salinity changes. According to previous studies, the pattern of correlation between stressors depends on the gradient of individual stressors (Fischer et al., 2012; Earhart et al., 2022; Mack et al., 2022; Segurado et al., 2022) and the target organisms (Corcoll et al., 2015).

**Fig. 4.**
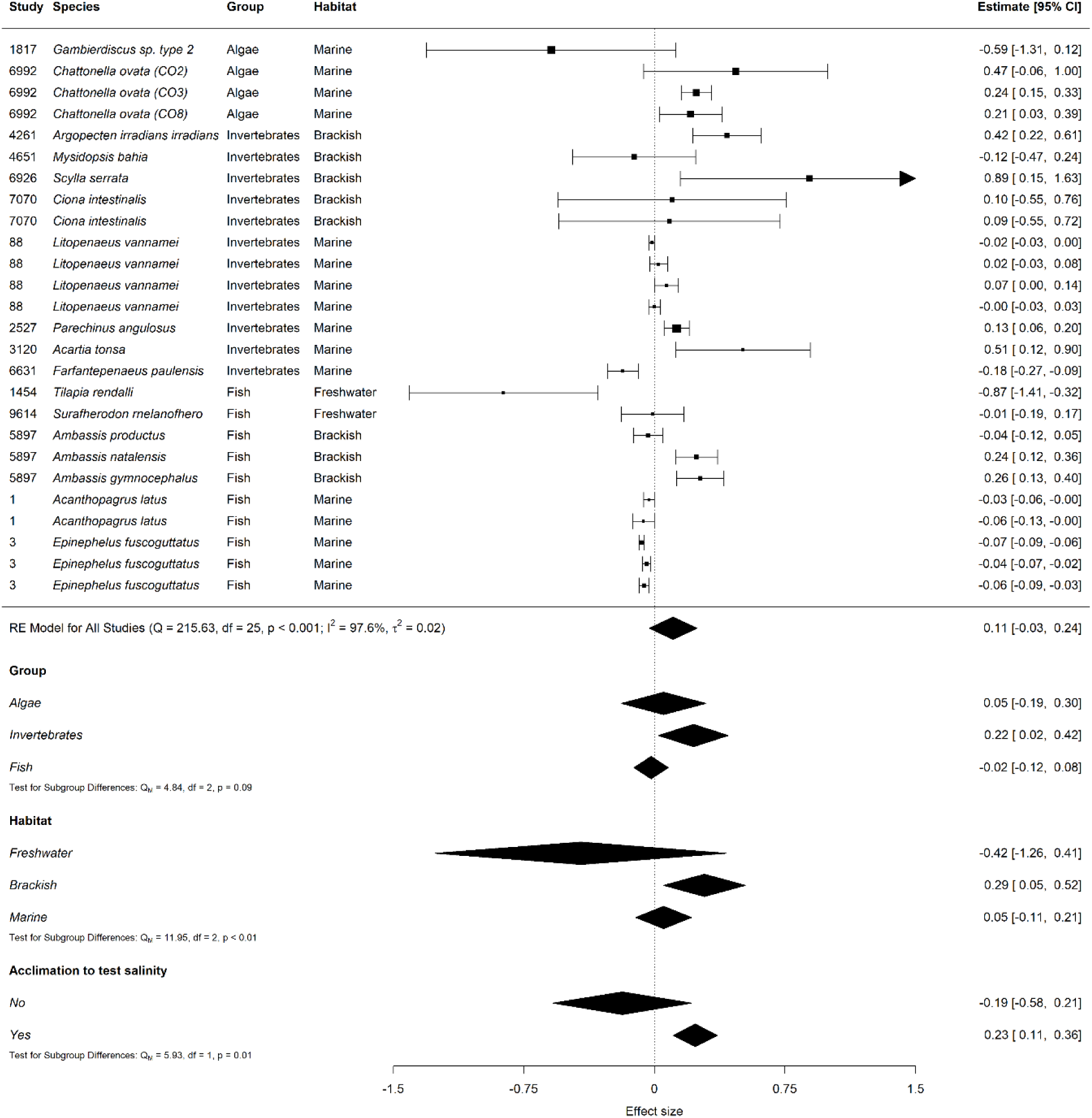
Effect of salinity on the thermal breadth. Mean (individual and pooled) effect size and 95% confidence interval were estimated from the model. Their values are listed on the right figure margin. The polygons at the bottom of the plot represent the mean effect size by different subgroups (organism group, habitat, and acclimation to test salinity). Significant effects are defined when the 95% confidence interval not overlapping with zero. Heterogeneity statistics of the model are shown by the values of the Cochran’s Q, its degree of freedom (df), *p*, I^2^, and τ^2^.

Salinity increases exerted opposite effects on the thermal tolerance of freshwater (negative) and of brackish/marine species (positive) (Fig. 3; Figs. S1-S3, Supporting Information). The thermal tolerance of freshwater species contracted, although not significantly, while the thermal tolerance of brackish and marine species significantly expanded with increasing salinity (Fig. 3; Figs. S1-S3, Supporting Information). This suggests different types of interactions between stressors in freshwater and marine systems, as demonstrated in previous meta-analyses (Crain et al., 2008; Harvey et al., 2013; Przeslawski et al., 2015; Jackson et al., 2016). Freshwaters and marine waters have dissimilar ion compositions with the former being dominated by Ca^2+^ and HCO_3_^-^ and the latter being dominated by Na^+^ and Cl^-^ (Wetzel, 2001). Moreover, the ions have different physiological functions (Charmantier et al., 2009). In freshwater animals, Na^+^ and Cl^-^ are major osmolytes responsible for maintaining the hyperosmotic state of their fluid (Bradley, 1987; Dietz, 1979; Evans, 2008; Wheatly and Gannon, 1995). Most freshwater animals are hyper-regulators that sustain higher ion concentrations in their blood or hemolymph compared to the external environment to achieve physiological homeostasis (Bradley, 2008). Most marine invertebrates are osmoconformers (Solan and Whiteley, 2016). For these organisms, it has been suggested that energy is not diverted to transport mechanisms in response to salinity changes as the osmolality of their internal medium fluctuates with the changing osmolality of the external medium (Rivera-Ingraham and Lignot, 2017). This contrasts with our result that salinity dilution contracted the thermal tolerance of marine species.

For freshwater organisms, experimental studies testing temperature effects at a range of different salinity levels (ideally, more than five temperature levels covering the width of the respective tolerance breadth of the tested species) are needed to address the central question of the present study, i.e., whether salinity gradients affect thermal tolerance of aquatic organisms. Even with such experimental designs, small shifts in temperature limits might be difficult to detect. It is possible that a modification of the critical thermal minimum/maximum framework applied for macroscopic organisms might be more suitable for addressing this question, but we are not aware of applications of this concept to microscopic organisms. In addition, a model linking cellular and molecular responses with responses at the organismal level might improve our understanding further.

### 3.2. Variation in the sensitivity of thermal tolerance to salinity gradients among organism groups

Our meta-analysis indicates insignificant variations of salinity effects on thermal tolerance among different groups of organisms (Fig. 5), contrasting with our hypothesis. The responsiveness of lower and upper temperature limits and the optimum temperature for different organism groups was: algae > invertebrates > fish (although not significant) (Fig. 5). It should be noted that the effect size in the present meta-analysis was estimated based on the slope of a linear relationship between thermal tolerance and salinity gradients. This method was selected because in most of the studies available, less than four salinity levels were included (Appendix 2). Mack et al. (2022) indicated that non-linear responses were more common for organisms at high trophic levels. In other words, the highest value of the critical thermal maximum is found at intermediate salinities. A non-linear relationship between the critical thermal maximum and salinity has been commonly found in fish (King and Sardella, 2017; Matern, 2001; Haney and Walsh, 2003; Sardella et al., 2008; Rodgers and Isaza, 2022). Such non-linear relationships might contribute to the dependence of co-tolerance on salinity levels. For example, for *Daphnia pulex*, Chen and Stillman (Chen and Stillman, 2012) reported cross-tolerance of temperature and salinity at intermediate salinity gradients and cross-susceptibility at extreme salinity variations.

**Fig. 5.**
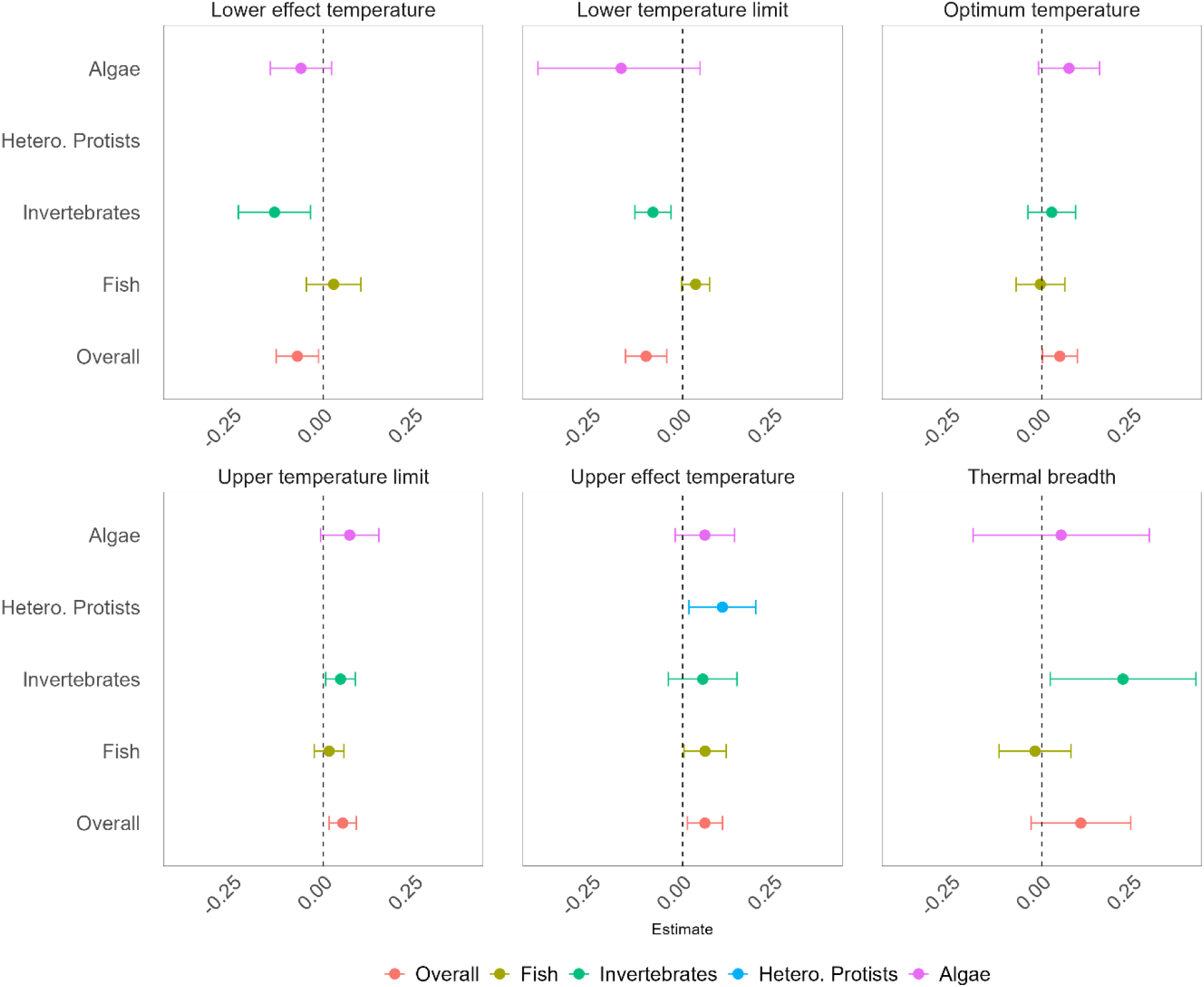
The effect size (mean; 95% confidence interval) of the influence of salinity changes on thermal indicators of algae, heterotrophic protists, invertebrates, and fish.

The more responsive thermal tolerance of algae to salinity gradients may be attributed to the higher sensitivity of algae to salinity gradients compared to other organism groups. Compared to larger multicellular organisms, unicellular organisms are more impacted by salinity as they have a limited capacity of osmotic buffering (Ishika et al., 2018). The size-dependent osmotic tolerance has been related to the different development of ionoregulatory capacity (Amiri et al., 2009) as larger organisms such as crustaceans and fish have developed osmoregulatory mechanisms (Charmantier et al., 2009; Evans and Claiborne, 2009). Further studies on the effect of multicellularity on the interactions of thermal and osmotic stress might provide more insights into the underlying mechanisms contributing to the variations among organism groups.

Only a few studies of algae (marine or freshwater, planktonic or benthic, macro- or microalgae) have applied a treatment matrix or other approach capable of detecting the interactive effects of both factors as illustrated in the present study. Bollen et al. (2016) found significant interactions between salinity and temperature effects upon photosynthetic maximum quantum yield for three (marine) kelp species, and for at least one species also upon photosynthetic electron transport rates, chlorophyll-a content, xanthophyll pool size, and antioxidant capacity. Bjaerke and Rueness (2004) did not find a significant interactive effect of temperature and salinity on the growth and survival of marine macroalga *Heterosiphonia japonica*, though some of their observations could possibly be interpreted as a narrowing of the thermal tolerance range at least at the highest desalination. In terms of microalgae studies, Xu et al. (2017) unravelled a pattern of decreasing temperature tolerance range for *Phaeocystis globosa* with suboptimal (decreasing) salinities. Similar observations were made for the marine diatom *Asteroplanus karianus* (Shikata et al., 2015). These latter studies, however, all investigated marine taxa and salinity concentrations below optimal level as a stressor, in contrast to a freshwater situation. Salinity stress with concentrations above the optimal level would be more relevant in the context of freshwater salinization. The publication of Le et al. (2023) was the only study we were able to locate that tested the interactive effects of temperature and salinity on the growth rate of six freshwater benthic diatom strains. In terms of publicly available data sets, the GlobTherm database contains information on the thermal tolerance of 274 red, brown, and green algal species, but none from the secondary endosymbiotic groups of microalgae playing important roles in many aquatic ecosystems (like diatoms, dinoflagellates; https://datadryad.org/stash/dataset/doi:10.5061/dryad.1cv08 last accessed 11.12.2023). More importantly, from the point of view of this study, GlobTherm contains no information on salinity conditions at which the individual thermal tolerances were determined.

Regardless of indicators, salinity changes consistently had a limited influence on the thermal tolerance of fish (Figs. 2-6). In fish, the intracellular and extracellular osmolality stabilises with a fluctuating osmolality of the external environment (Griffith, 2017; Urbina and Glover, 2015). Walker et al. (2020) pointed out that there is an isosmotic point in fish, i.e., an internal osmolality that is equal to the osmolality of the external environment, below which less energy is required for osmoregulation. Moreover, fish species in our database are mostly euryhaline and in some cases, even anadromous (e.g., salmon). These species have adapted to changes in their osmotic environment and are less likely to be challenged by salinity stress. Previous research indicated that other factors that can influence thermal tolerance in fish, especially the upper thermal limit, are acclimation temperature, photoperiod, and duration (Hines et al., 2019). The influence of photoperiod on thermal tolerance also varies between species. Usually, longer photoperiods enhance thermal tolerance (Healy and Schulte, 2012; Terpin et al., 1976); however, artificially long photoperiods can also act as a stressor, and inhibit thermal tolerance (Hines et al., 2019; Newman et al., 2015). In the study of Hines et al. (2019), the upper temperature tolerance decreased during a 400-day experiment from ∼29°C to ∼26°C in all salinity treatments (Hines et al., 2019). In such long experiments, the effect could be additionally affected by the substantial changes in body size (Recsetar et al., 2012). Effects reported from the field (e.g., Morgan et al., 2019) are therefore likely related to additional stressors such as energy limitation, predation, or pathogen pressure.

In contrast to algae and fish, the influence of salinity gradients on thermal tolerance in invertebrates has been more intensively studied. But still, most of the studies have been mainly conducted on marine species. More intensive research on marine species allows for conducting meta-analyses. These studies yielded inconsistent patterns of interactions between temperature and salinity. The meta-analysis in the study of Crain et al. (2008) indicated antagonistic interactions as the most common pattern, while Przeslawski et al. (2015) suggested synergistic interactions. Such differences might be attributed to the different life stages and phyla considered in these two studies (Lange and Marshall, 2017), and/or to the species-specific response of osmoregulation capacities to temperature (Torres et al., 2021). Combined effects of temperature and salinity have been evaluated on some freshwater invertebrate species (Kumlu et al., 2010; Venancio et al., 2023). For instance, Venancio et al. (2023) suggested synergistic effects of these factors on the survival of *Daphnia longispina*.

### 3.3. Other confounding factors

#### Experimental conditions

Acclimation to test salinity significantly enhanced the thermal tolerance of aquatic organisms (Fig. 3; Figs. S1-S3, Supporting Information). Acclimation strongly affected temperature and salinity tolerance in laboratory experiments (Botella-Cruz et al., 2016; Fernandes et al., 2023), where – after a short acclimation period – the lower temperature limit was above the limit that was observed under natural (*in situ*) conditions (White et al., 2015). The importance of acclimation to test salinity was also unravelled in our study as shown by the significant difference in the effect size on the upper temperature limit between the acclimated and unacclimated organisms. This result agrees with the acclimation-enhanced tolerance reported in a number of previous studies (Loureiro et al., 2015; Kumlu et al., 2010). For instance, acclimation of fish from various habitats to high temperatures led to significantly higher tolerance towards high temperatures (e.g., Davis et al., 2019; Matern, 2001), while the acclimation to lower temperatures increased tolerance to low temperatures (King and Sardella 2017; Schofield et al., 2009). Acclimation temperature therefore had a strong effect on the thermal tolerance, and the profound differences between the studies might have concealed a general salinity effect. However, this moderator was not considered in the model because this information is not given in many studies.

##### Relevance of parasite infections in evaluating stressor effects

An important group of organisms in the context of studies focusing on stress responses are parasites. Parasites can directly and/or indirectly modulate ecological interactions, affecting, for instance, mobility, habitat selection, foraging, reproduction, longevity, and morphology of infected hosts (Fenton and Rands, 2006; Forbes, 1993; Hurd et al., 2001; Miura et al., 2006). More importantly, parasites may alter host tolerance to stressors such as fluctuating salinity and increasing temperature, sometimes with surprising outcomes (Sures et al., 2023). For instance, the acanthocephalan *Polymorphus minutus* enhanced the salinity tolerance of its host *Gammarus roeselii* (Piscart et al., 2007). Studies dealing with environmental stressors should, therefore, account for parasitic infections in their study organisms, or else they are at risk of biased outcomes to some degree, as already seen in the ecotoxicological context (Grabner et al., 2023). As studies analysed in the current work have not taken the influence of parasitism into account, variations in the data could, for instance, be due to unrecognized infections, and the here-presented results should rather be interpreted in the context of uninfected individuals or, at least of those not strongly affected by their parasites.

Although the final dataset did not contain any records on parasites, they would be an important target group for the analyses presented here due to their overall complexity. Parasites can be protists or metazoans, external (ectoparasites) or internal (endoparasites), have simple (involving one host) or complex (involving multiple hosts) life cycles, be host generalist or host specialist, and have differing life stages (Dobson et al., 2008; Lafferty, 2012). All these strategies can influence the outcome of parasite-stressor interactions, resulting in parasite species, populations, or their life stages responding differently to salinity and temperature changes (Sures et al., 2023).

For example, ectoparasites are directly exposed to the external environment. Hence, their tolerance to temperature and salinity changes, which might be lower than that of the host, is directly affected by environmental changes. This is exemplified by aquaculture practices in which salt or freshwater baths are employed to treat hosts infected with freshwater and marine parasites (Buchmann, 2022). On the other hand, endoparasites are protected from the external environment by the host, therefore experiencing stressors somewhat indirectly. For instance, host osmoregulation mitigates external changes in salinity, maintaining an environment suited for the development of endoparasites (Möller, 1978). Nevertheless, endoparasites may be exposed to the external environment at some stage of their lifecycle. Spores (microparasites), eggs (macroparasites), and free-living larval stages (micro- and macroparasites) are de facto challenged by environmental changes (Sures et al., 2023). Hence, most multiple stressors studies on endoparasites focus on their free-living stages. However, these stages are rather resistant to temperature and salinity changes, and the survival of endoparasites is often indirectly constrained by the host’s tolerance to stressors, but not by the direct impact of stressors (Möller, 1978; Rogowski and Stockwell, 2006).

Despite their high relevance and multifaceted life strategies, it seems that parasites have often been seen as an additional stressor on their hosts rather than a separate entity on which multiple-stressor effects should be assessed. Consequently, studies dealing solely with the combined effects of temperature and salinity on freshwater parasites are scarce. In the future, more resources should be allocated to investigate the impact of environmental stressors on both parasites and their hosts.

## 4. Conclusions

The influence of salinity changes on the thermal breadth of aquatic organisms has been investigated mostly on brackish and marine species. Studies on freshwater species are required to achieve a more comprehensive evaluation on the influence of salinization. Our meta-analysis indicates salinity changes-elevated effects of thermal stress, supporting the hypothesis that energetically expensive osmoregulation in response to salinity changes might lead to effects on the thermal tolerance of aquatic organisms. The effect of salinity on the thermal tolerance differed, but not, significantly among groups of organisms with the following order of responsiveness: algae > invertebrates > fish. Furthermore, the infection status of the host should be considered in further analyses as the thermal tolerance of the host might be affected by parasitism.

## Acknowledgements

This study was performed within the Collaborative Research Centre 1439 RESIST (Multilevel Response to Stressor Increase and Decrease in Stream Ecosystems; www.sfb-resist.de) funded by the Deutsche Forschungsgemeinschaft (DFG, German Research Foundation; CRC 1439/1, project number: 426547801).

## Notes

### Competing Interest Statement

The authors have declared no competing interest.

