## Appendix 1 for "Influence of salinity on the thermal tolerance of aquatic organisms"

Appendix 1. Database construction

A database of thermal tolerance at various salinity levels was built by retrieving the relevant variables (e.g. optimum temperature, temperature limits, thermal breadth, and incipient lethal temperature) from peer-reviewed publications available through the Web of Science and Scopus. The literature survey was conducted on 18$^{th}$ January 2024 with the following search terms and combinations thereof: toleran* OR sensitiv* OR optimal temperatur* OR optimum temperatur* OR critical temperatur* OR ctmax OR incipient lethal temperature OR thermal limit OR ctmin OR critical thermal minimum OR critical thermal maximum OR thermal breadth; AND temperatur* OR thermal* OR heat OR warm*; AND salin* OR salt* OR conductiv* OR osmo* OR total dissolved solids OR sodium chloride; AND effect* OR impact* OR respons* OR influence* OR modulat* OR change OR affect OR modif*; AND invertebrat* OR diatom* OR fish* OR pisc* OR macroinvert* OR algae* OR parasit*; AND aquati* OR freshw* OR river* OR stream* OR lake OR coast* OR lagoo* OR marin* OR brack* OR intertid* OR estuar* OR water OR creek* OR pond*. After checking for duplicates, the literature survey resulted in 9,624 unique records.

The records were screened following a stepwise approach. In the first step, the machine learning tool ASReview (Active learning for Systematic Reviews) was applied for a systematic screening using 44 relevant and 45 irrelevant records as a training dataset. Relevant records were defined as studies on thermal tolerance at various salinity conditions and studies on the effects of temperature and salinity. Irrelevant records included; field studies or field study-based modelling research; studies on community, meta-community, or ecosystem responses; studies on non-biological factors only; studies on irrelevant organisms (terrestrial organisms and partially terrestrial vertebrates); and studies on irrelevant subjects (food or food processing; extracellular or intracellular biochemistry; extraction or purification; uptake, absorption or bioaccumulation; waste treatment or pollutant removal; device or methods; and palaeo- or history records). ASReview screening provide a ranking order from the most relevant to the most irrelevant based on the training. A preliminary screening with a dataset including 500 randomly-selected records excluded around 45% of the dataset. Therefore, records ranked after 4000 from the whole dataset screening were manually screened with the above exclusion criteria, and this process was stopped when 50 continuous records were assigned irrelevant, assuming that the records ranked below are irrelevant.

After the ASReview abstract screening, 6,384 records (66 %) remained for the second screening step, i.e. manual abstract screening by domain experts (screeners). A validation dataset of 50 records was taken from these records and distributed to seven screeners. Besides the exclusion criteria employed during the ASReview screening, we proceeded with the manual exclusion of review studies, studies that missed temperature or salinity effects or both, and studies based on the influence of temperature and/or salinity on other stressors. The results obtained from all screeners were highly consistent, with more than 90% overlap of keepers. Most importantly, irrelevant records were correctly identified and excluded. The entire set of 6,384 records was randomly assigned to the screeners. Relevant records remaining after the manual abstract screening (1,345 records; 21%) were randomly distributed to four screeners for intensive screening. In this step, publications were screened based on the following criteria: 1) Records provide at least one indicator of thermal tolerance (Fig. 1; Table S1, Supporting Information). Indicators of thermal tolerance include those listed in Figure 1 and ultimate temperatures: ultimate lower incipient lethal temperature (i.e. the lower incipient lethal temperature that does not decrease with decreasing acclimation temperatures) and ultimate upper incipient lethal temperature (i.e. the upper incipient lethal temperature that does not increase with increasing acclimation temperature), and the ultimate critical thermal minimum and maximum (with similar interpretation regarding the influence of acclimation temperature); 2) Effects of temperature and salinity were simultaneously investigated; 3) Data were reported for at least three salinity levels; 4) Data were obtained from laboratory experiments or modelling research based on laboratory experiments in controlled conditions in which other factors besides temperature and salinity were consistently maintained among treatments; and 5) Data were based on measured responses at the individual level in aquatic-life phases of the organism. We excluded studies in which thermal tolerance was determined under irrelevant environmental conditions, for example, air exposure.
