## Supporting information for "Influence of salinity on the thermal tolerance of aquatic organisms"

Table S1. Data availability for various indicators of thermal tolerance

| Indicator | Definition | Data availability |
|---|---|---|
| Optimum temperature | Temperature at which performance (i.e. growth) is maximal | 29 |
| Optimum survival temperature | Temperature at which survival is maximal | 13 |
| Preferred temperature | The range of temperatures in which animals congregate or spend the most time in a gradient or free-choice situation | 4 |
| Critical thermal minimum | Lower level of the range within which physiological failure (i.e. cessation of movement) does not occur | 22 |
| Critical thermal maximum | Upper level of the range within which physiological failure (i.e. cessation of movement) does not occur | 33 |
| Physiological thermal breadth | Difference between critical thermal minimum and critical thermal maximum (generalist) or been 50% CTmin and 50% CTmax (specialist) | 13 |
| Lower lethal temperature | Lower level of the 0% mortality range (i.e. within the range all test individuals survive) | 12 |
| Upper lethal temperature | Upper level of the 0% mortality range (i.e. within the range all test individuals survive) | 12 |
| Survival thermal breadth | Difference between lower lethal temperature and upper lethal temperature | 8 |
| Lower incipient lethal temperature | Lower level of the range outside which less than 50% of test individuals survive | 28 |

| | | |
|---|---|---|
| Upper incipient lethal temperature | Upper level of the range outside which less than 50% of test individuals survive | 35 |
| Lower incipient physiological temperature | Lower level at which performance halves of the maximum | 21 |
| Upper incipient physiological temperature | Upper level at which performance halves of the maximum | 19 |
| Lower 100% mortality temperature | Lower level of the 100% mortality range (i.e. outside the range all test individuals die) | 11 |
| Upper 100% mortality temperature | Upper level of the 100% mortality range (i.e. outside the range all test individuals die) | 11 |
| Ultimate critical thermal minimum | Critical thermal minimum that does not decrease with decreasing acclimation temperature | 1 |
| Ultimate critical thermal maximum | Critical thermal maximum that does not increase with increasing acclimation temperature | 1 |
| Ultimate lower incipient lethal temperature | Temperature bellow which no decrease in the incipient lethal temperature is accomplished by further decreases in acclimation temperature | 1 |
| Ultimate upper incipient lethal temperature | Temperature beyond which no increase in the incipient lethal temperature is accomplished by further increases in acclimation temperature | 2 |

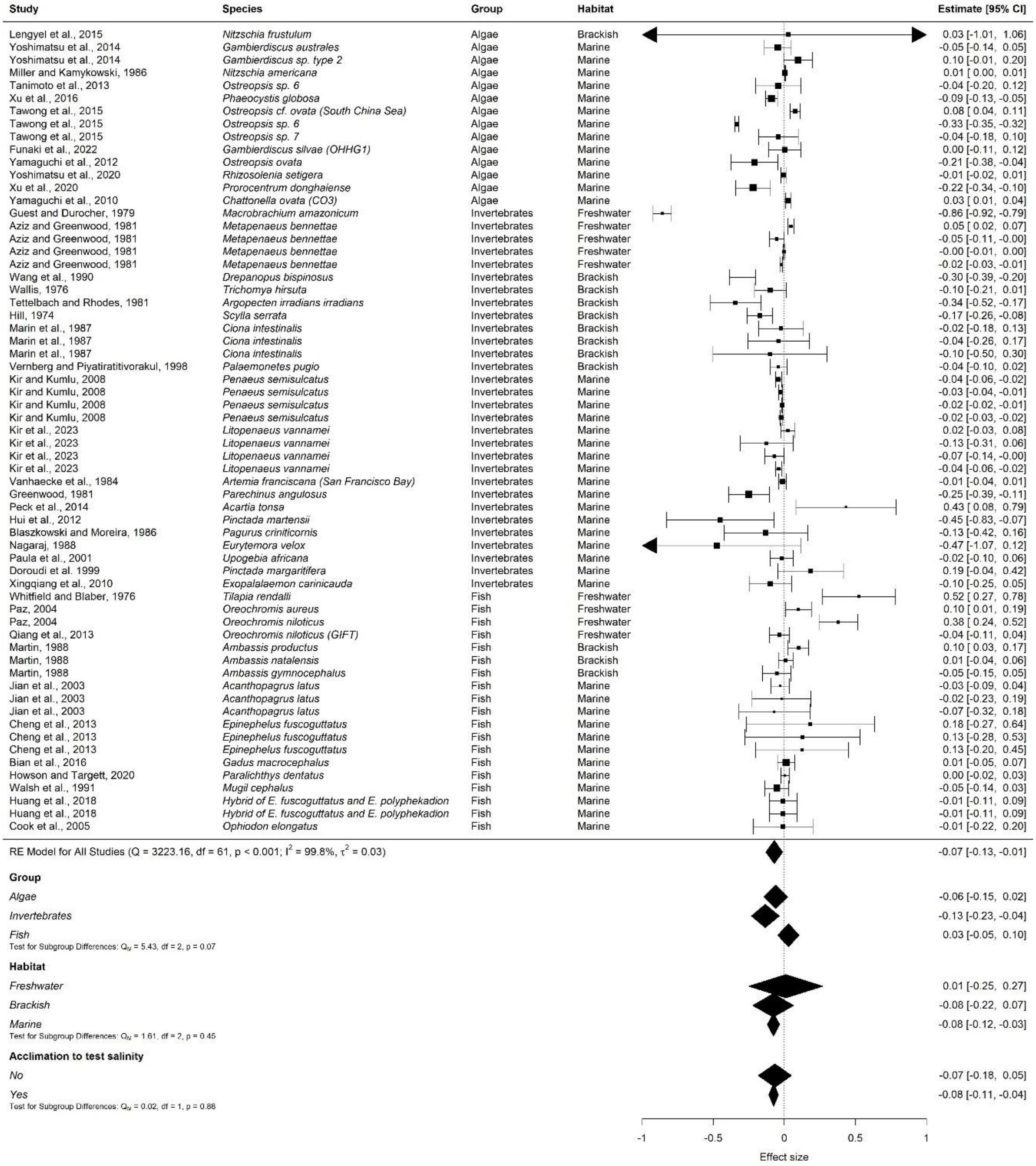


Fig. S1. Effect of salinity on the lower effect temperature. Mean (individual and pooled) effect size and 95% confidence interval were estimated from the model. Their values were listed on the right side of the figure. The polygons at the bottom of the plot represent the mean effect size by different subgroups (organism group, habitat and acclimation to test salinity). Significant effects are defined as the 95% confidence interval not overlapping with zero.

Heterogeneity statistics of the model are shown by the values of the Cochran's Q, its degree of freedom (df), $p$, $I^2$, and $\tau^2$.

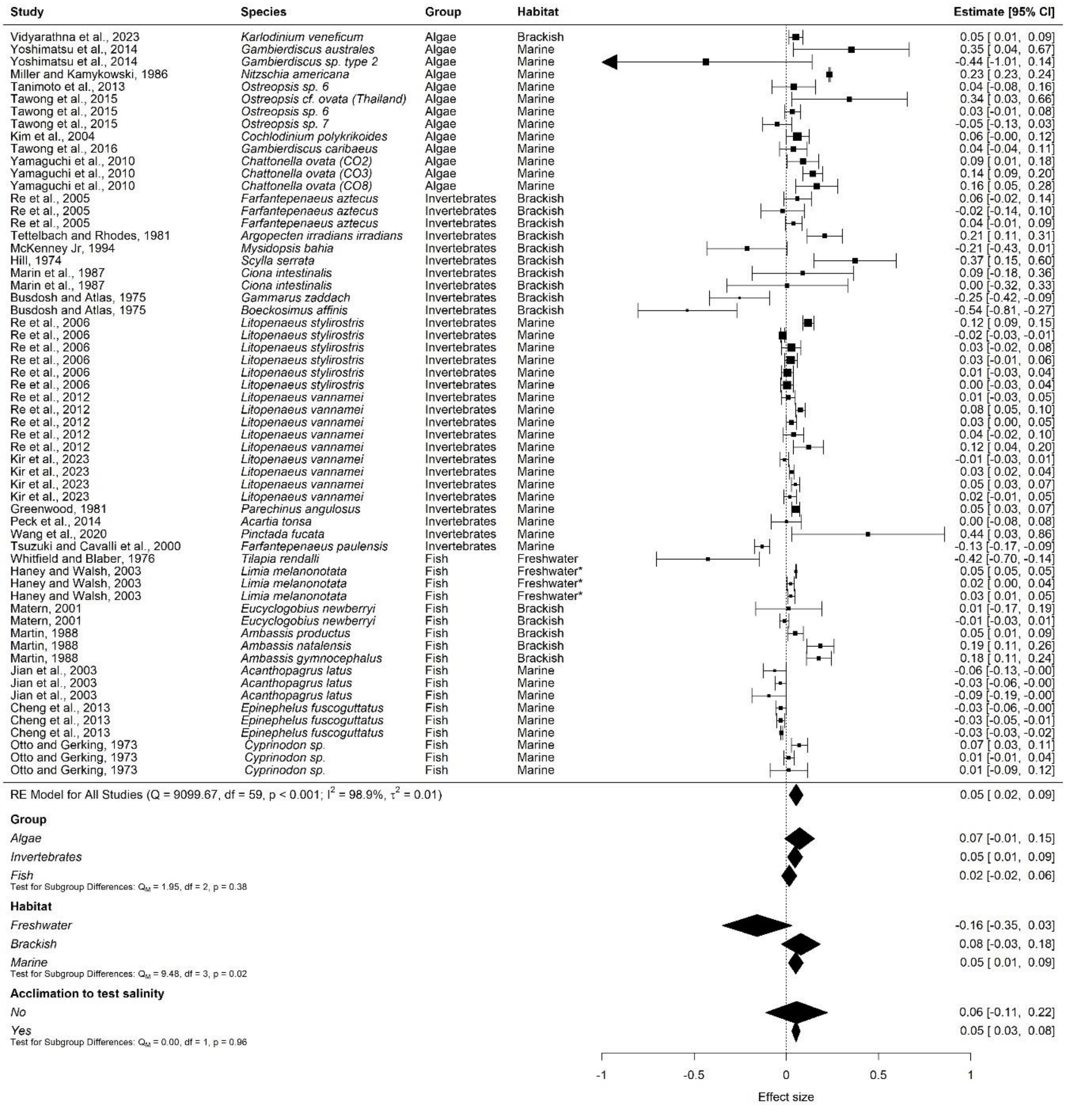


Fig. S2. Effect of salinity on the upper temperature limit. Mean (individual and pooled) effect size and 95% confidence interval were estimated from the model. Their values were listed on the right side of the figure. The polygons at the bottom of the plot represent the mean effect size by different subgroups (organism group, habitat and acclimation to test salinity).

Significant effects are defined as the 95% confidence interval not overlapping with zero. Heterogeneity statistics of the model are shown by the values of the Cochran's Q, its degree of freedom (df), *p*, $I^2$, and $\tau^2$.

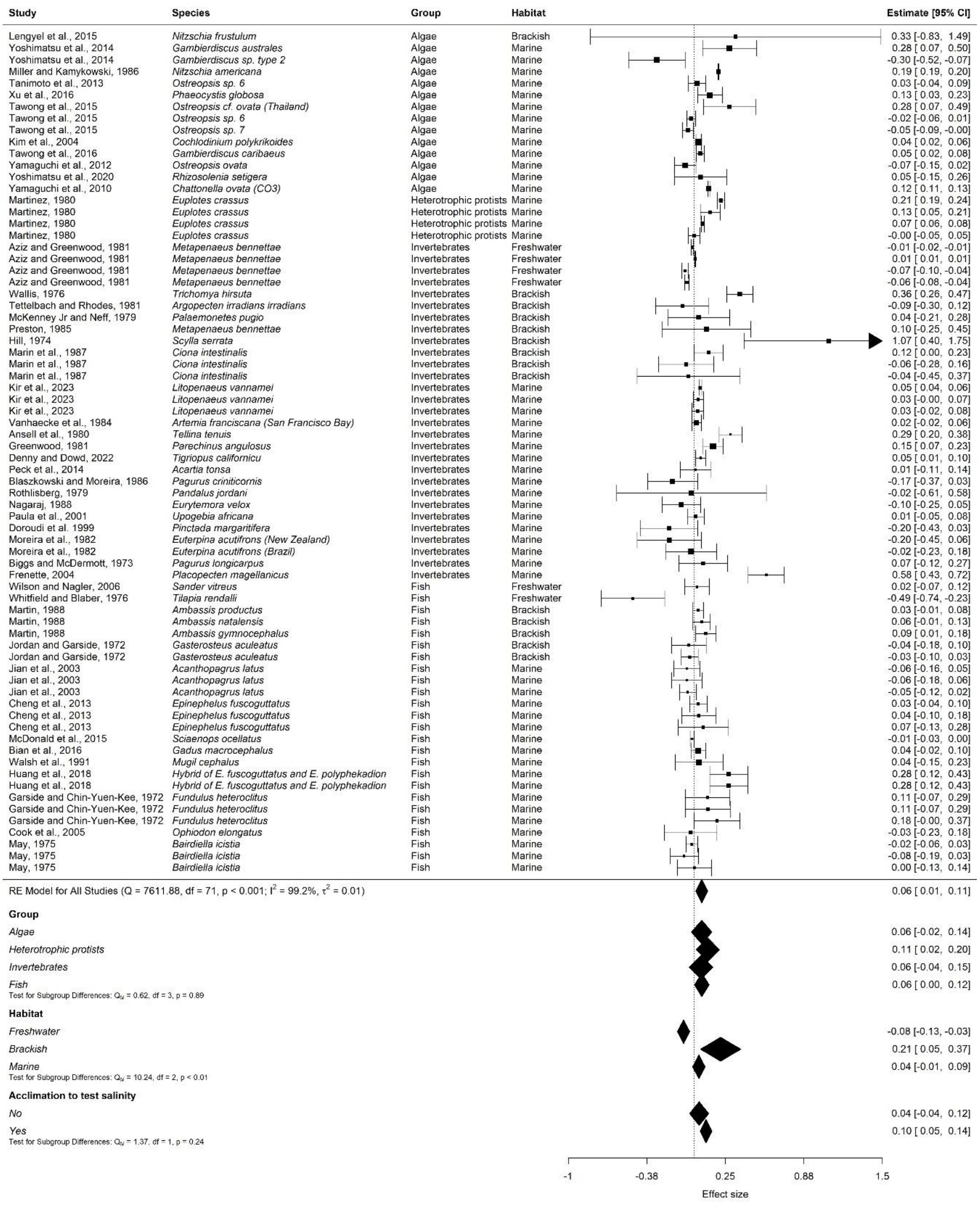


Fig. S3. Effect of salinity on the upper effect temperature. Mean (individual and pooled) effect size and 95% confidence interval were estimated from the model. Their values were listed on the right side of the figure. The polygons at the bottom of the plot represent the mean effect size by different subgroups (organism group, habitat and acclimation to test salinity).

Significant effects are defined as the 95% confidence interval not overlapping with zero. Heterogeneity statistics of the model are shown by the values of the Cochran's Q, its degree of freedom (df), $p$, $I^2$, and $\tau^2$.

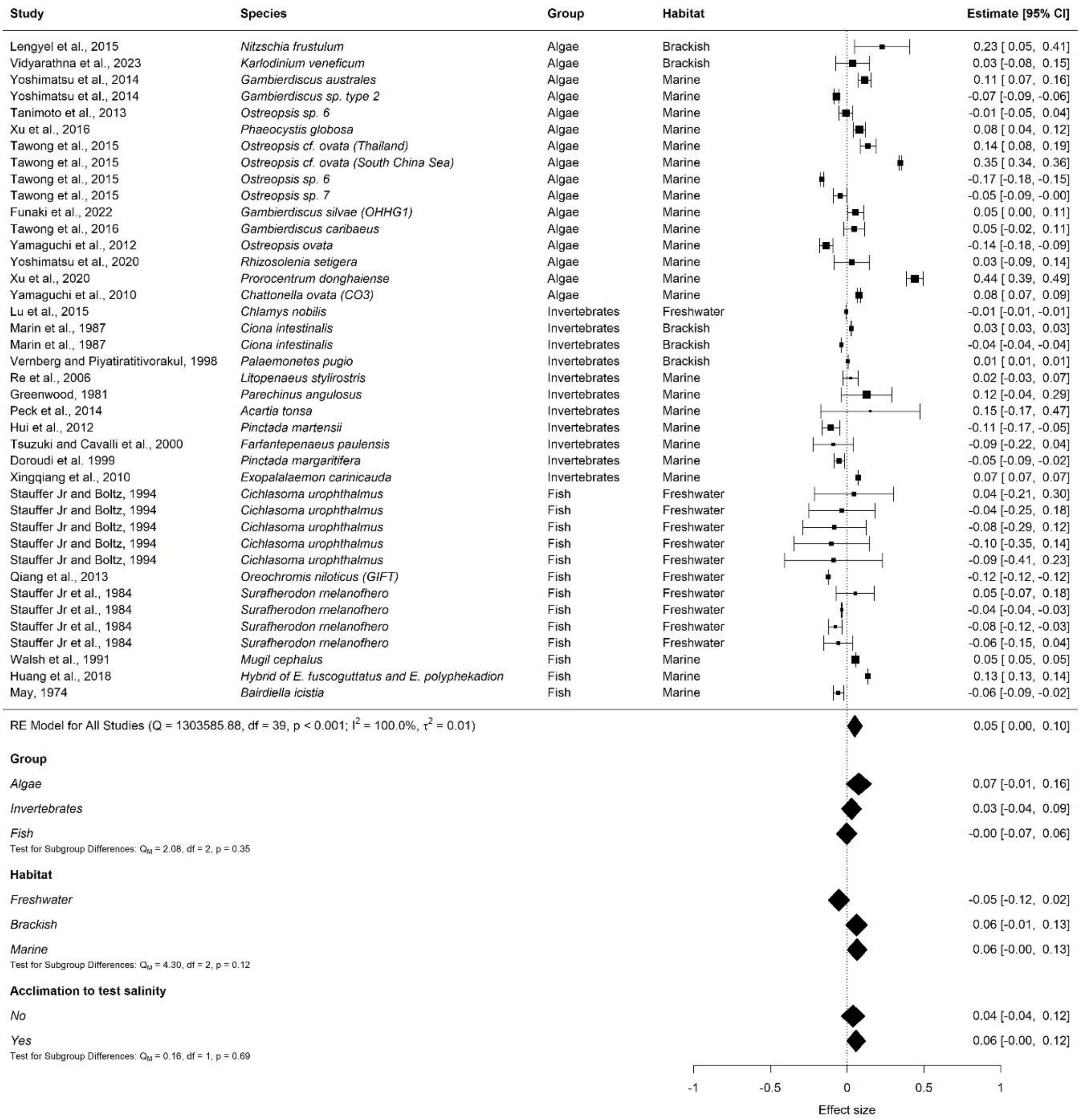


Fig. S4. Effect of salinity on the optimum temperature. Mean (individual and pooled) effect size and 95% confidence interval were estimated from the model. Their values were listed on the right side of the figure. The polygons at the bottom of the plot represent the mean effect size by different subgroups (organism group, habitat and acclimation to test salinity). Significant effects are defined as the 95% confidence interval not overlapping with zero. Heterogeneity statistics of the model are shown by the values of the Cochran's Q, its degree of freedom (df), $p$, $I^2$, and $\tau^2$.
